# Frequency-Dependent Bioimpedance Signatures of Ocular Tissues in Intact Ex Vivo Eyes Under Simulated Surgical Conditions

**DOI:** 10.64898/2026.05.14.725195

**Authors:** Bita Behziz, Matthew Nepo, Yasamin Sadat Mousavimotlagh, Tsu-Chin Tsao, Aya Barzelay

## Abstract

**Purpose:** To characterize the frequency-dependent bioimpedance properties of major ocular tissues in intact ex vivo porcine eyes under simulated surgical conditions and evaluate tissue separability at discrete frequencies.

**Methods:** Bioimpedance spectra were acquired from sclera, corneal epithelium, iris, lens, vitreous, and retina in intact ex vivo porcine eyes using a two-electrode probe and a precision LCR meter over 5 kHz to 1 MHz. Measurements were obtained under balanced salt solution and ophthalmic viscosurgical device conditions. Probe–tissue contact was verified by microscope visualization and optical coherence tomography. Tissue separability at 5, 50, 100, and 900 kHz was evaluated using global and pairwise statistical comparisons, effect sizes, and ROC-based separability metrics. Robotic-stabilized and handheld measurements were also compared.

**Results:** Ocular tissues demonstrated distinct, frequency-dependent impedance magnitude distributions. Across sampled frequencies, 60% to 80% of tissue pairs showed significant differences after multiplicity correction. Median pairwise effect sizes ranged from Cohen’s d = 0.48 at 5 kHz to 1.04 to 1.06 at 50 to 100 kHz. Median ROC-based separability was 0.91 at 5 kHz and 0.76 to 0.77 at 50 to 900 kHz. Robotic-stabilized measurements showed lower variance than handheld measurements, although tissue-specific impedance ranges and frequency-dependent trends were preserved across acquisition modes.

**Conclusions:** Major ocular tissues exhibit reproducible, frequency-dependent bioimpedance signatures in intact ex vivo eyes under simulated surgical preparation. These findings establish a physiologically relevant ocular impedance reference dataset and support bioimpedance as a complementary modality for tissue differentiation in ophthalmic microsurgery.

## Introduction

Intraocular microsurgery requires submillimeter precision near fragile retinal and anterior-segment structures. However, intraoperative decision-making relies primarily on visual feedback, with limited or absent tactile sensation, particularly in teleoperated robotic workflows. Unintended contact with delicate tissues can result in iatrogenic injury, including retinal trauma or anterior-segment damage ^1,2^. The absence of intrinsic haptic feedback in robotic-assisted surgery may further increase this risk ^3^. A sensing modality capable of identifying tissue at the instrument tip could therefore provide immediate feedback and reduce the likelihood and severity of unintended tissue interaction.

Bioimpedance sensing is a promising complementary modality because biological tissues exhibit frequency-dependent electrical properties that reflect differences in composition, structure, and extracellular environment. Bioimpedance has been widely used to discriminate tissue types and pathological states across multiple organ systems, including lung, breast, liver, and brain tissue ^4-8^. In contrast to pathology-focused applications, the present study aims to generate a library of impedance signatures for major ocular tissues encountered during cataract and vitreoretinal surgery and to quantify tissue-to-tissue separability under simulated surgical conditions.

In ophthalmology, prior studies have demonstrated the feasibility of impedance-based proximity sensing and tissue identification. For example, Schoevaerdts et al. demonstrated bioimpedance-based proximity sensing in vitreoretinal microsurgery ^9^, and Aghajani Pedram et al. developed a tissue identification framework integrating impedance sensing with machine learning for cataract surgery ^10^. However, most ocular bioimpedance studies have been conducted using isolated tissues or simplified in vitro preparations. While these approaches are valuable for initial validation, they do not preserve intact ocular geometry, surrounding media, or tissue adjacency, all of which influence the electrical conduction pathway between electrodes.

Measurements acquired in intact ex vivo eyes better preserve anatomical boundaries and boundary conditions compared with isolated tissues, while still allowing controlled experimental access. In clinical settings, intraocular media are continuously altered by balanced salt solution (BSS) infusion and fluid exchange, which further influence impedance measurements. In the present study, eyes were intermittently irrigated with BSS and prepared with ophthalmic viscosurgical device (OVD) to approximate a wet-field surgical environment without active regulation of intraocular pressure.

In this work, we characterize the frequency-dependent bioimpedance behavior of major ocular tissues in intact ex vivo porcine eyes under simulated surgical preparation and quantify tissue-to-tissue separability at discrete sensing frequencies. Impedance measurements were obtained from sclera, corneal epithelium, iris, lens, vitreous, and retina, with probe–tissue contact verified using microscope visualization and optical coherence tomography (OCT). To improve placement repeatability during reference library construction, the probe was integrated with the Intraocular Robotic Interventional and Surgical System (IRISS) ^11,12^, which served as a stabilizing platform. To evaluate translational robustness beyond robotic stabilization, handheld measurements acquired by an experienced ophthalmic surgeon were also analyzed.

Prior work in robotic-assisted impedance sensing has demonstrated the value of stabilization for proximity detection and tissue characterization ^9-12^. However, handheld operation introduces probe–tissue interface variability, including changes in contact pressure, micro-motion, tissue deformation, and hydration state, all of which can influence bioimpedance measurements across frequencies ^13-18^. In this study, handheld measurements obtained by an experienced ophthalmic surgeon were therefore analyzed as a practical stress-test of measurement robustness relative to robotic-stabilized acquisition.

Agreement between acquisition modes is quantified using intraclass correlation and Bland–Altman analysis, and conditions under which impedance distributions overlap are explicitly identified. Together, these results establish an intact-eye ocular bioimpedance reference dataset and define the degree of frequency-dependent separability achievable across major ocular tissues under surgically relevant boundary conditions.

## Methods

To simulate surgical access, conventional corneal incisions and pars plana sclerotomies were created to enable probe insertion into the anterior and posterior segments. During experimentation, the ocular surface and surgical field were intermittently irrigated with balanced salt solution (BSS) to maintain hydration and approximate a clinical wet-field environment. Ophthalmic viscosurgical device (OVD) was introduced into the anterior chamber to simulate intraoperative conditions. Intraocular pressure was not actively regulated.

To enable localized impedance measurement at the instrument tip, a two-electrode integrated probe design was implemented. The probe consisted of two conductors that were electrically insulated from one another except at a small, exposed region (approximately 1 mm) at the distal tip. This configuration constrained the measured electrical pathway to the tissue in direct contact with the probe tip, ensuring that impedance measurements primarily reflected local tissue properties^10^.

The mechanical structure of the probe was designed to be compatible with standard surgical instrumentation. A standard 18-gauge (1.2 mm) stainless steel needle was attached to a 1 mL syringe. The metallic needle served as one electrode, while a second insulated conductor was inserted through the syringe barrel and aligned with the needle tip. Both conductors were connected to the measurement system, forming a localized two-electrode sensing interface at the probe tip. The same probe was used across all measurements to ensure consistency.

The probe was carefully inserted through the pars plana using the IRISS controller to facilitate precise data collection. Bioimpedance measurements were acquired using a precision LCR meter (E4980A, Keysight Technologies, Santa Rosa, CA, USA), which served as both the excitation source and measurement device. During acquisition, the probe was mounted to the Intraocular Robotic Interventional and Surgical System (IRISS), which served as a positioning and stabilization platform. The probe was advanced using the IRISS controller until contact with the target tissue was achieved. Upon contact, the tissue completed the electrical pathway between the two electrodes, allowing impedance magnitude to be measured from the resulting current response to the applied excitation signal. Figure 1 illustrates the experimental setup, including probe integration with the robotic system and the measurement workflow.

**Figure 1:**
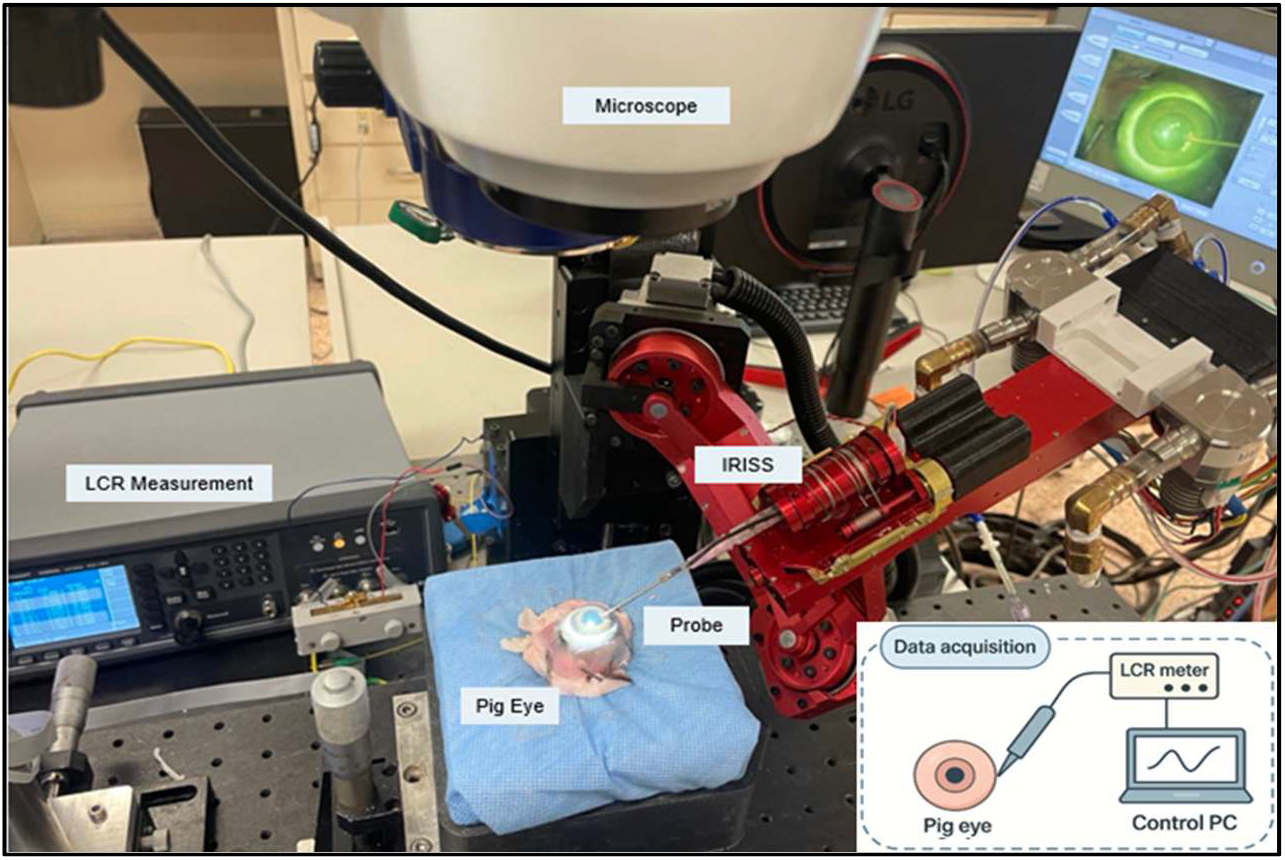
Experimental and measurement setup—Probe assembly and installation on the IRISS robotic manipulator, recording on ocular samples. During measurements of the cornea and sclera, the ocular surface was continuously rehydrated with balanced salt solution (BSS) using a syringe (not pictured) to maintain physiological moisture conditions. The bottom-right inset shows a schematic of the data-acquisition workflow, including the probe–tissue interface, LCR meter, and PC.

During data collection, the robotic arm was teleoperated until the probe contacted the target tissue. Probe–tissue contact was confirmed using both microscopic visualization and OCT. Figure 2 presents representative microscopic and corresponding OCT images of the probe in contact with sclera, epithelial cornea, iris, lens, vitreous, and retina. These tissues were selected because they are major ocular tissues encountered during cataract and vitreoretinal surgery and therefore provide a clinically relevant basis for impedance characterization. It is important to note that, for consistency and to reduce the influence of localized bioimpedance effects within sensitive tissues, all measurements of individual tissues within each eye were performed 1-2 mm apart from the previous location of measurement. Additionally, measurement of the iris occurred at the midpoint between the iris root and the sphincter at the iris stroma. Both corneal epithelium and endothelium were measured. Epithelial measurements were acquired under robotic positioning during closed-globe preparation. Endothelial measurements were acquired under an open-sky, manual positioning condition due to robotic workspace limitations. Because endothelial measurements were not collected under the same closed-globe robotic protocol, they are reported separately (Figure 6C) and were not included in the discrete-frequency separability analysis. Before each tissue was measured in an individual eye, air and BSS were measured as a baseline comparison. If air and BSS reference measurements were within a preset acceptance range, the probe was considered clean and stable for the next tissue measurement. If the expected value was not achieved, the probe was cleaned and air and BSS were measured again. This process was repeated until the probe had been sufficiently cleaned. While impedance measurements were conducted under closed-globe conditions, for improved visualization and clarity for the readers of this paper, the microscopic and OCT images illustrating the probe’s position within the vitreous and retina are shown under open-sky conditions (figure 2, E and F).

**Figure 2:**
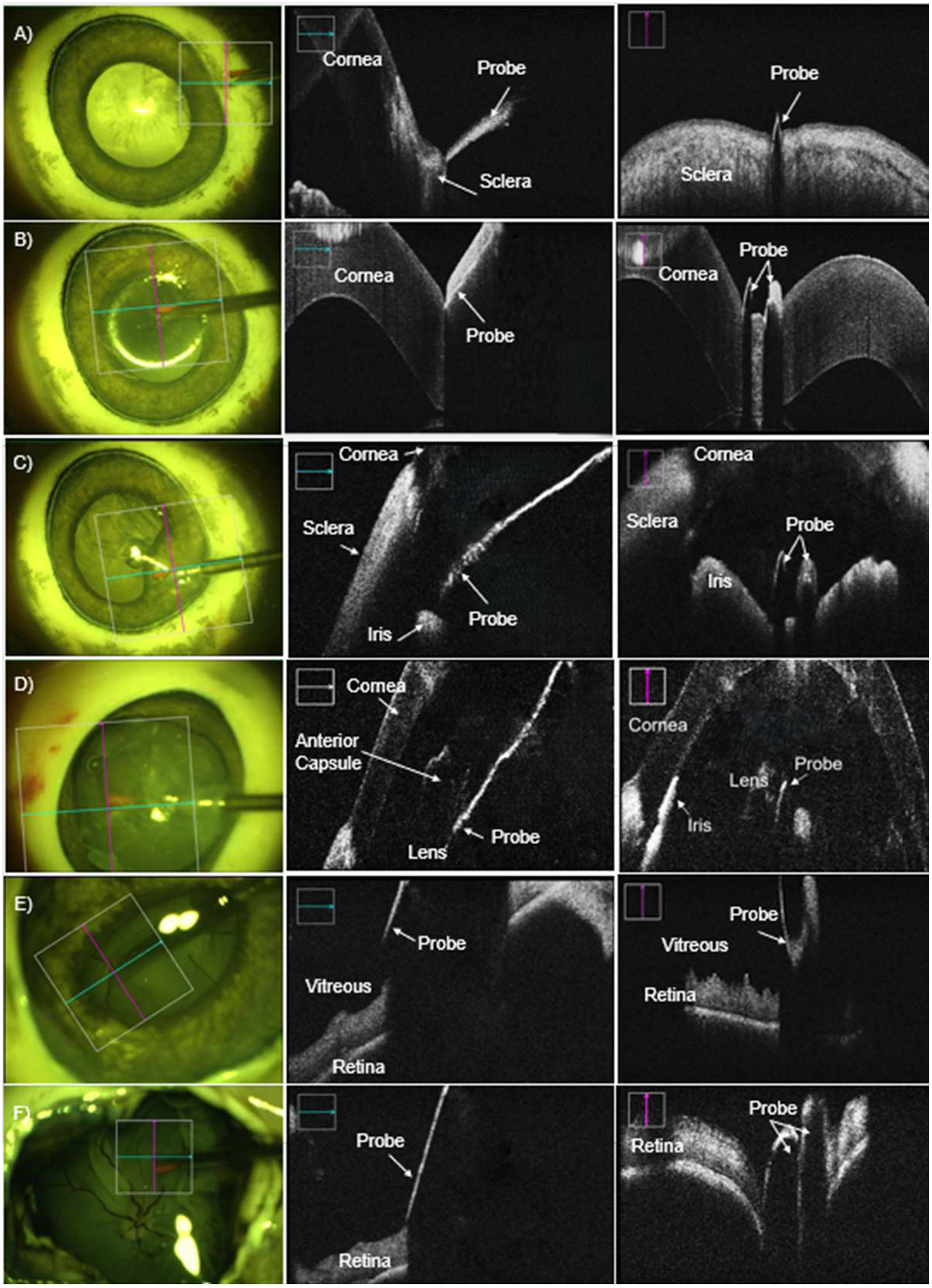
Real-time microscopic view and OCT scans (Longitudinal (blue) and transverse (red)) illustrating the contact of the bio-impedance probe with ocular tissues under closed-globe condition for A) sclera, B) cornea, C) iris, D) lens, and under open-sky conditions for E) vitreous, and F) retina.

For each medium (BSS, vitreous, OVD, and a composite mixture), three impedance sweeps were acquired using the same frequency-sweep protocol used for tissue measurements shown on figure 6. Each measurement was performed upon the same eye in a different, 1-2 mm from the previous location of measurement, to maintain consistency regarding localized bioimpedance effects. Room temperature was used for all measurements; corneal and scleral hydration was maintained by intermittent BSS irrigation (30-s intervals; ∼5-s wait before recording). Intraocular pressure was not measured or actively regulated; irrigation was used to maintain a wet-field, hydrated surface rather than to control physiological pressure. Preliminary feasibility data was collected using 4 porcine eyes; the reported reference libraries were constructed from an additional 12 porcine eyes. The ages of the eyes used range from approximately 5 hours to 3 days post-mortem.

Prior bioimpedance studies have identified low- and mid-range frequencies as informative for tissue discrimination^14,15^. In this study, discrete-frequency analyses were performed at 5, 50, 100, and 900 kHz to construct the reference impedance library. These frequencies were selected to sample low-, mid-, and high-frequency behavior across the spectra while capturing regions of strong contrast and partial plateau behavior observed in the tissue impedance curves. This plateau behavior is observable in both the means and the standard deviations of each tissue’s impedance values, as illustrated in Table 4.

Statistical analyses were performed to quantify inter-tissue differences at each sampled frequency shown in figure 5. We report global group tests (one-way ANOVA or Welch ANOVA, depending on variance equality), followed by multiplicity-corrected post-hoc pairwise comparisons (Tukey HSD or Games–Howell) and effect sizes.

### Animal tissue statement

Porcine eyes were obtained post-mortems from an external source for research use. No live animals were used in this study. All procedures adhered to the ARVO Statement for the Use of Animals in Ophthalmic and Vision Research.

### Statistical analysis

The primary outcomes were tissue impedance magnitude as a function of frequency, pairwise tissue separability at discrete frequencies, and agreement between robotic-stabilized and handheld acquisition modes. Multiple measurements were acquired per tissue within each eye at sites separated by 1–2 mm. For statistical analysis, measurements were averaged within each tissue per eye, and separability analyses were performed using these per-eye averaged values to account for within-eye correlation. Impedance magnitude was analyzed at discrete frequencies and across full sweeps to characterize frequency-dependent behavior and quantify inter-tissue separability. For discrete-frequency comparisons among tissues (and media where applicable), we first assessed overall group differences using one-way ANOVA (or Welch ANOVA when variances were unequal), followed by post-hoc pairwise comparisons (Tukey HSD or Games–Howell as appropriate) with multiplicity correction. In addition to p-values, we report effect sizes (η^2^ for global tests; Cohen’s d for pairwise contrasts) and 95% confidence intervals.

To quantify practical discriminability beyond global statistical significance, we computed the pairwise receiver operating characteristic area under the curve (ROC–AUC) at each discrete frequency. Because ROC–AUC depends on label assignment, we report a direction-invariant separability metric defined as AUC_sep = max(AUC, 1–AUC). In addition, for each tissue pair and frequency, we determined a single-threshold decision rule that maximized balanced accuracy. These analyses provide quantitative measures of tissue separability and distributional overlap without training a multivariate classifier.

For robotic versus manual agreement, we quantified agreement using intraclass correlation coefficients (ICC), Bland– Altman analysis, and paired comparisons of matched tissue measurements.

Post-mortem age effects were treated as exploratory: we report descriptive trends and (when sample size supports it) a mixed-effects model with eye as a random effect and age as a fixed effect, rather than hypothesis tests on interpolated values.

### Dataset and sample characterization

Impedance measurements were obtained from sclera, corneal epithelium, iris, lens, vitreous, and retina in intact ex vivo porcine eyes. Lens measurements were performed with residual intraocular fluid present in the anterior segment; however, because eyes were ex vivo and not under physiologic perfusion or intraocular pressure regulation, this fluid cannot be assumed to represent in vivo aqueous humor. Eyes were stored at 4°C without chemical fixation and allowed to equilibrate to room temperature prior to acquisition. Time post-mortem ranged from approximately 5 hours to 3 days (Table 1). No live animals were used in this study; eyes were harvested off-site and provided for experimental use.

**Table 1:** Characteristics of Ex-Vivo Porcine Eyes Used for Bioimpedance Measurements. time postmortem presented in hours.

| Age | <24 Hr | 24-48 Hr | 48-72 Hr | 72 Hr< |
| --- | --- | --- | --- | --- |
| Number of Eyes | 5 | 2 | 6 | 2 |

The following dataset was generated under standardized simulated surgical preparation conditions. Conventional corneal incision and pars plana sclerotomies were created to allow probe access. The ocular surface and surgical field were intermittently irrigated with balanced salt solution (BSS) to maintain hydration and approximate a clinical wet-field environment; intraocular pressure was not actively regulated. Ophthalmic viscosurgical device (OVD) was introduced into the anterior chamber to simulate phacoemulsification conditions.

Measurements were acquired while tissues remained in their intact anatomical configuration. Unlike isolated in vitro tissue preparations, the intact-eye configuration preserves tissue adjacency and surrounding media that influence the electrical conduction pathway between electrodes. However, these ex vivo conditions do not replicate physiological perfusion or intraocular pressure regulation present in vivo. Continuous balanced salt solution infusion and washout effects characteristic of clinical surgery were not implemented.

For each tissue within an eye, impedance measurements were repeated at multiple sites separated by 1–2 mm to reduce localized site dependence and to better approximate within-tissue variability under practical acquisition conditions. Media baselines (air and BSS) were measured periodically to verify probe stability and cleanliness prior to tissue acquisition.

### Post-Mortem Age Analysis

To assess potential variability associated with post-mortem interval, impedance distributions were examined across the 5-hour to 3-day window represented in the dataset (Figure 3).

**Figure 3:**
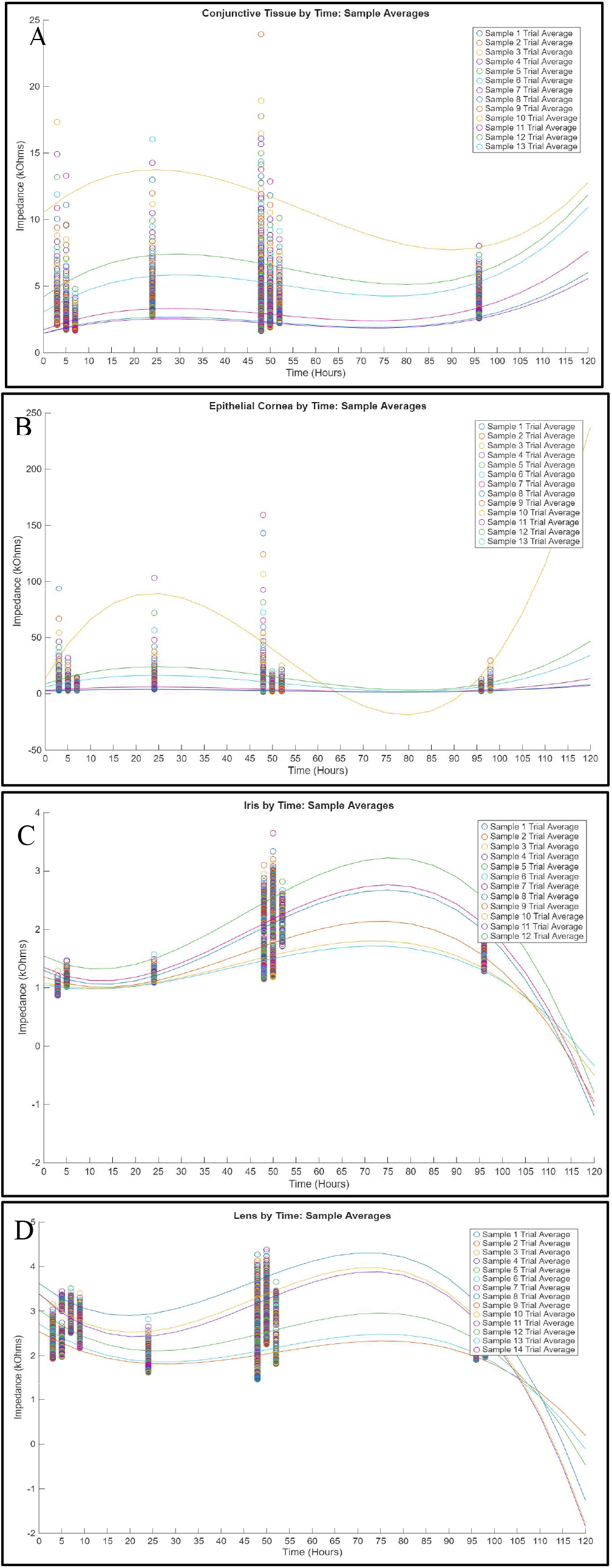
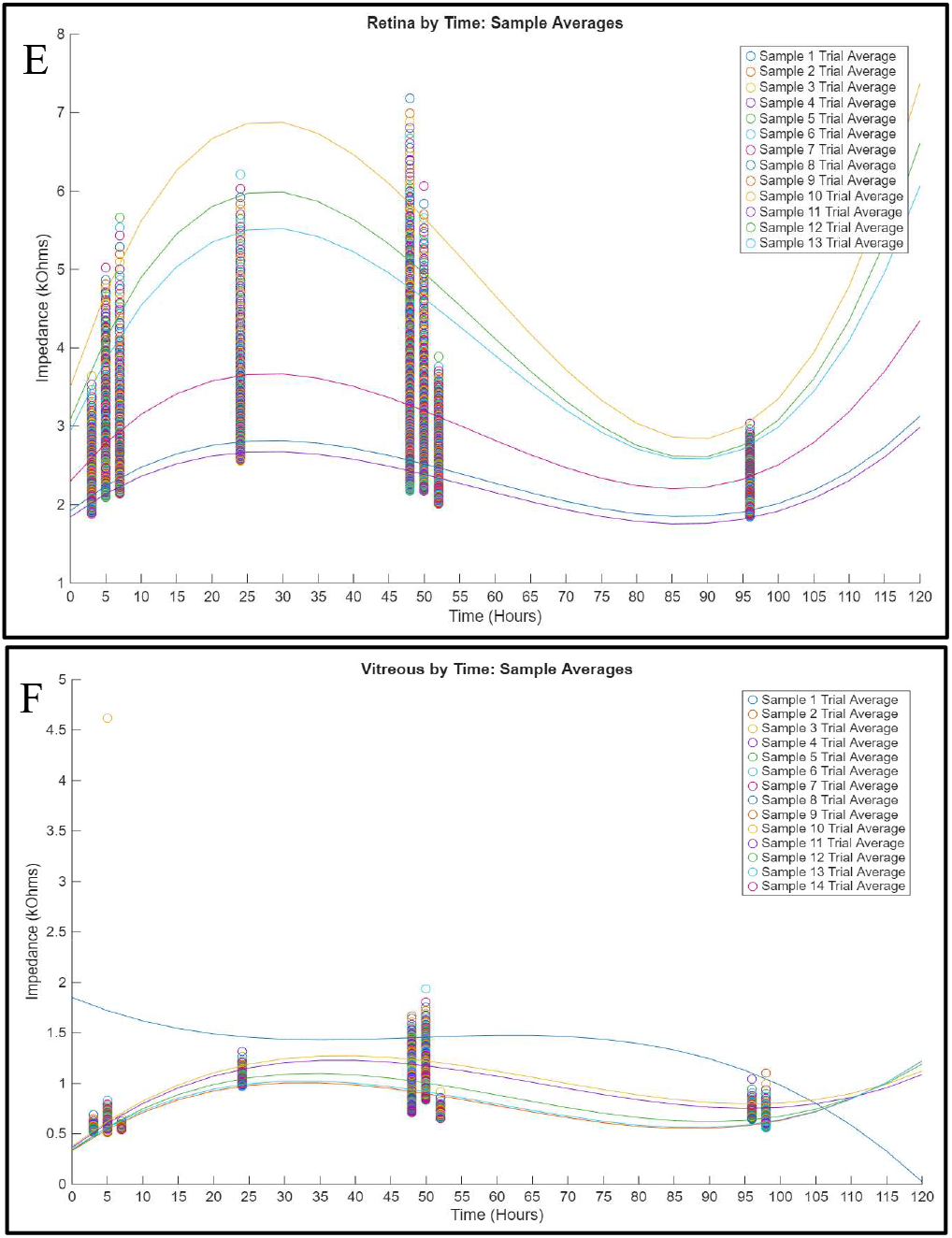
Interpolated Data for Age Stratification (Measured Bioimpedance Range vs. Age in Hours) for A) Sclera, B) Epithelial Cornea, C) Iris, D) Lens, E) Retina, and F) Vitreous.

Because sample sizes within individual age strata were limited and unevenly distributed, post-mortem effects are reported descriptively rather than inferentially. No hypothesis tests were performed on interpolated values, and the present dataset does not provide sufficient statistical power to establish age-independence within this post-mortem window. For visualization purposes only, smooth curves in Figure 3 were generated using cubic spline interpolation applied to mean impedance values at each post-mortem time point; interpolated values were not used for statistical testing or inference. While variability is observed across eyes, no consistent monotonic trend is evident within the sampled window. However, the present dataset is not powered to detect subtle age-dependent effects.

Accordingly, post-mortem interval is treated as a potential source of variability in the impedance library rather than as a negligible factor. Future studies with balanced sampling across tightly controlled post-mortem intervals will be required to rigorously quantify age-related impedance changes.

## Results

To evaluate whether ocular tissues exhibit frequency-dependent impedance characteristics under simulated surgical conditions, impedance measurements were acquired from intact ex vivo porcine eyes immersed in balanced salt solution (BSS) and ophthalmic viscosurgical device (OVD) environments. Preliminary data was collected from four eye samples across a frequency range of 5 kHz to 1MHz by the Keysight LCR meter. Figure 4 evaluates tissue-to-background contrast within relevant surgical media (BSS, OVD or gel, and composite intraocular fluid), demonstrating that tissue contact produces impedance values distinct from surrounding surgical media under simulated intraoperative conditions. These panels establish the probe’s ability to detect transitions from fluid or viscoelastic media to solid tissue, but do not test tissue-to-tissue separability. Accordingly, Figure 4 is interpreted strictly as evidence of tissue–medium contrast rather than inter-tissue classification. Each tissue was measured within its native physiological medium, and balanced salt solution (BSS) irrigation was performed at 30-second intervals to maintain hydration.

**Figure 4:**
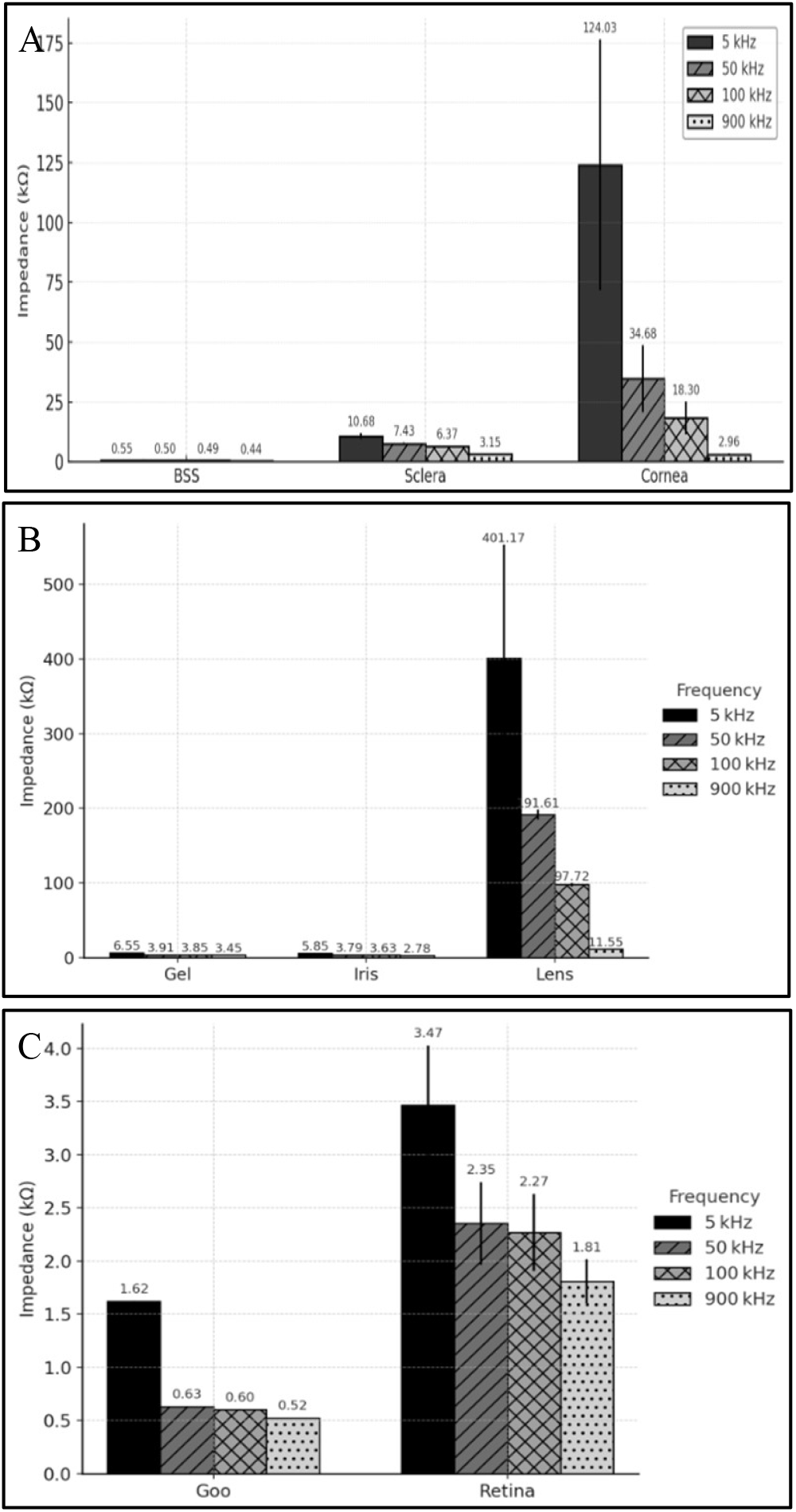
Tissue–medium contrast under simulated surgical media. Impedance (kΩ) is shown for (A) cornea and sclera relative to BSS, (B) iris and lens relative to OVD/gel, and (C) retina relative to composite intraocular media (“Goo”). These panels demonstrate tissue–medium contrast under simulated surgical conditions. They are not intended to establish tissue-to-tissue separability.

Once tissue–background contrast was established in Figure 4, impedance data across tissues were evaluated at discrete frequencies to assess overall differences between tissue groups. Figure 5 displays impedance distributions at selected frequencies (5, 50, 100, and 900 kHz). The reported p-value in each panel represents the global significance across all tissue groups at that frequency (ANOVA/Welch ANOVA). A significant global p-value indicates that at least one tissue differs from the others but does not establish pairwise tissue separability. Therefore, tissue-to-tissue separability was quantified using multiplicity-corrected post-hoc pairwise comparisons with effect sizes (Table 2) and by computing pairwise ROC–AUC and threshold-based balanced accuracy at each frequency (Table 3).

**Figure 5:**
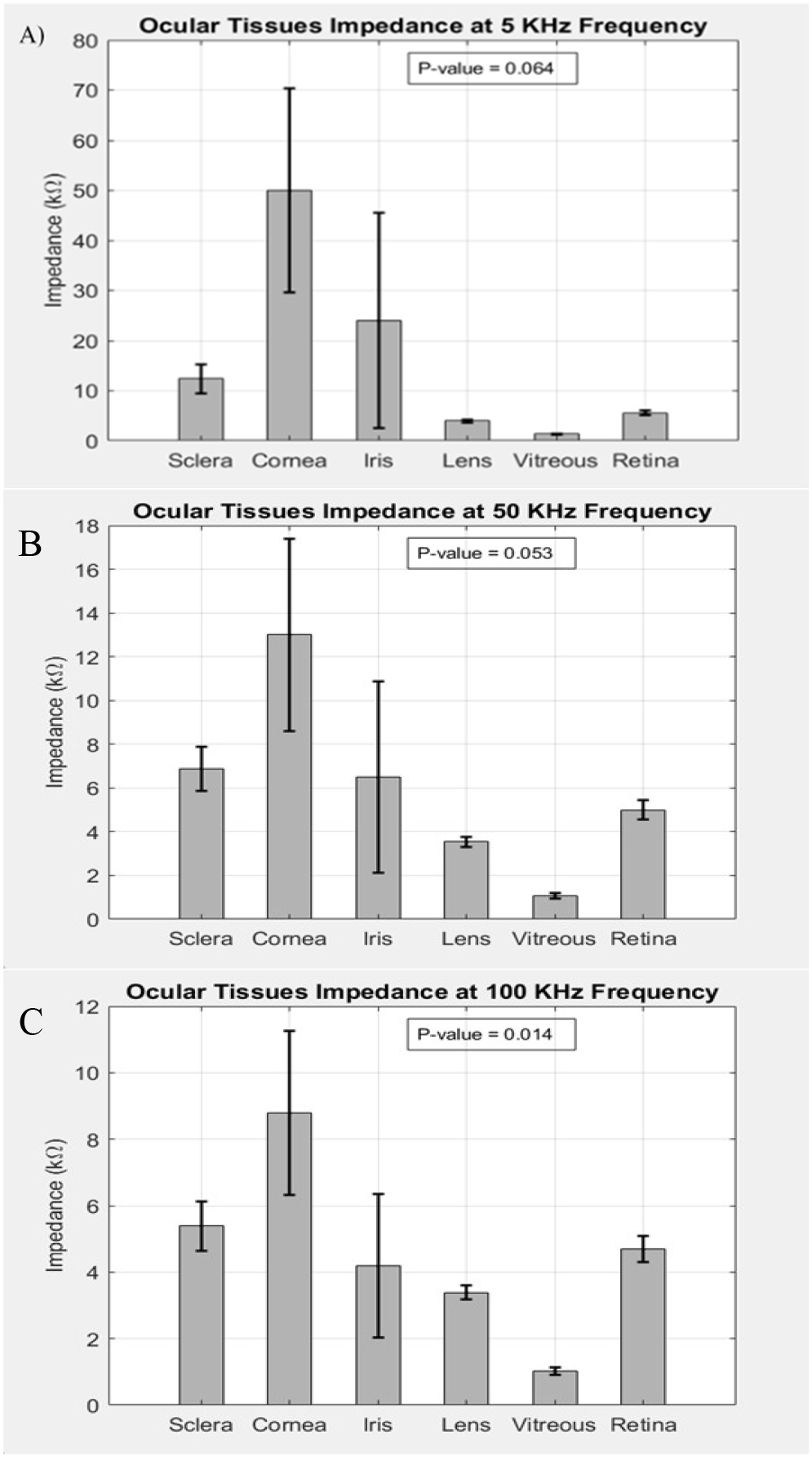
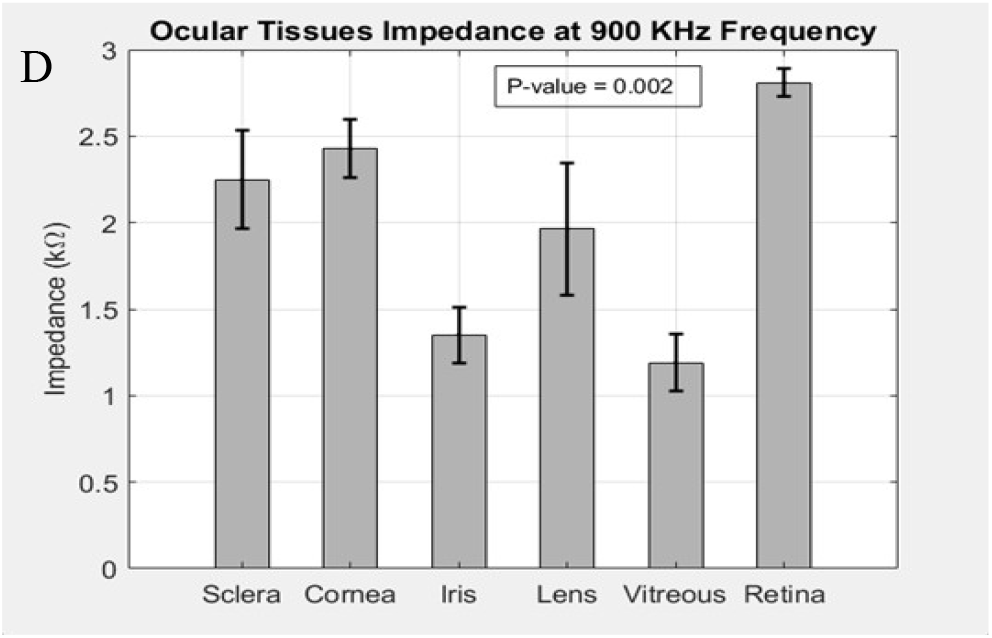
Discrete-frequency impedance distributions across ocular tissues. Impedance magnitude (kΩ) distributions are shown for each tissue at A) 5, B)50, C)100, and D)900 kHz. The p-value in each panel indicates the global statistical significance across all tissue groups at that frequency (ANOVA or Welch ANOVA, as appropriate). A significant global p-value indicates that at least one tissue differs from the others but does not establish pairwise separability. Pairwise tissue separability is evaluated using multiplicity-corrected post-hoc comparisons (Table 2) and ROC–AUC with threshold-based balanced accuracy (Table 3).

**Table 2:** Summary of pairwise statistical separability between ocular tissues across discrete frequencies. Multiplicity-corrected post-hoc comparisons were performed following global ANOVA/Welch ANOVA to evaluate impedance differences between tissue pairs at 5, 50, 100, and 900 kHz. The table reports the percentage of tissue pairs demonstrating statistically significant differences (adjusted p < 0.05), the median effect size (Cohen’s d), and the range of observed effect sizes across all pairwise comparisons at each frequency. These summary metrics quantify the overall magnitude and consistency of impedance-based tissue separability across frequencies. Full pairwise comparison results, including adjusted p-values and confidence intervals, are provided in Supplementary Table S1.

| Frequency (kHz) | % significant pairs ($p < 0.05$) | Median effect size (Cohen's $d$) | Effect size range |
| --- | --- | --- | --- |
| 5 kHz | 60 | 0.475471587 | -1.94–3.47 |
| 50 kHz | 80 | 1.059791579 | -1.28–4.38 |
| 100 kHz | 80 | 1.038981878 | -1.41–4.39 |
| 900 kHz | 80 | 0.990572583 | -1.64–4.74 |

**Table 3:**
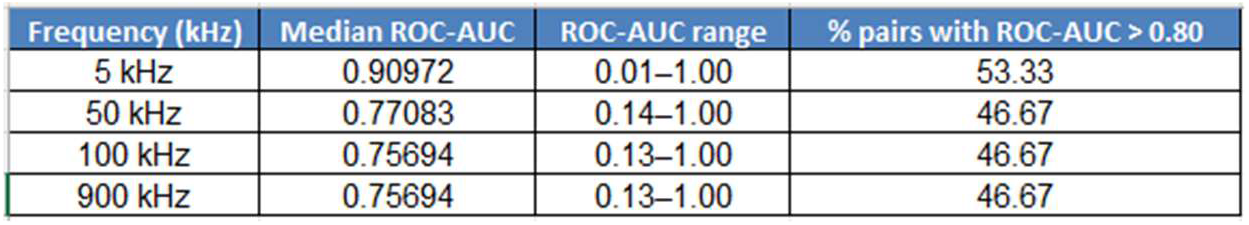
Summary of ROC–AUC–based separability between ocular tissues across discrete frequencies. Pairwise receiver operating characteristic area under the curve (ROC–AUC) and threshold-based balanced accuracy were computed to quantify impedance-based separability between tissue pairs at 5, 50, 100, and 900 kHz. The table reports median ROC–AUC values, range of ROC–AUC values, and the percentage of tissue pairs demonstrating strong separability (ROC– AUC > 0.80) at each frequency. ROC–AUC provides a threshold-independent measure of distributional separability, while balanced accuracy reflects achievable discrimination using simple threshold-based classification. Full pairwise ROC–AUC and balanced accuracy results are provided in Supplementary Table S2.

| Frequency (kHz) | Median ROC-AUC | ROC-AUC range | % pairs with ROC-AUC $> 0.80$ |
| --- | --- | --- | --- |
| 5 kHz | 0.90972 | 0.01–1.00 | 53.33 |
| 50 kHz | 0.77083 | 0.14–1.00 | 46.67 |
| 100 kHz | 0.75694 | 0.13–1.00 | 46.67 |
| 900 kHz | 0.75694 | 0.13–1.00 | 46.67 |

Separability varied systematically as a function of both tissue pair and sampling frequency (Tables 2 and 3). Multiplicity-corrected post-hoc comparisons confirmed statistically significant impedance differences across the majority of tissue pairs, with effect sizes increasing from moderate at 5 kHz to large at 50–100 kHz. ROC–AUC analysis demonstrated moderate-to-strong discrimination across frequencies, with several tissue contrasts achieving strong separability (ROC–AUC > 0.80), while others remained moderate (0.60–0.80), particularly in cases where impedance distributions overlapped.

Importantly, separability performance was not uniform across frequencies. Mid-range frequencies (50–100 kHz) provided the most consistent discrimination across tissue pairs, reflected by larger effect sizes, higher ROC–AUC values, and a greater proportion of statistically significant comparisons. In contrast, both low-frequency (5 kHz) and high-frequency (900 kHz) measurements exhibited increased variability and reduced contrast for specific tissue pairs. These findings indicate that impedance-based tissue differentiation is inherently frequency-dependent and that optimal sensing performance requires frequency selection tailored to the tissue contrasts of interest rather than reliance on a single universal frequency.

To characterize frequency-dependent behavior beyond discrete sampling points, impedance magnitude was plotted across the full frequency sweep for the sclera tissue, epithelial cornea, iris, lens, vitreous, and retina, while intact in their surgical environment, which is OVD and BSS (Figure 6).

**Figure 6:**
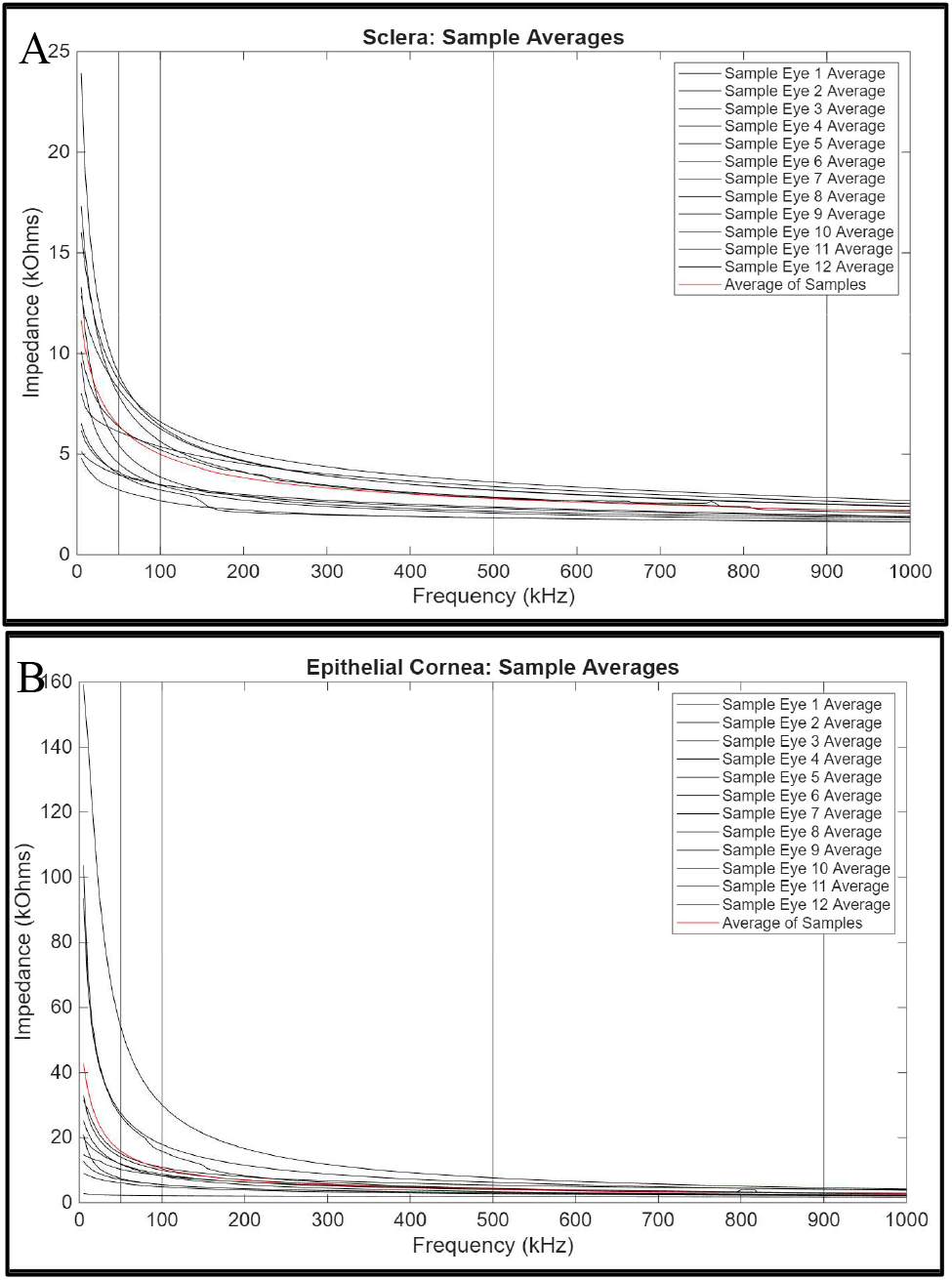
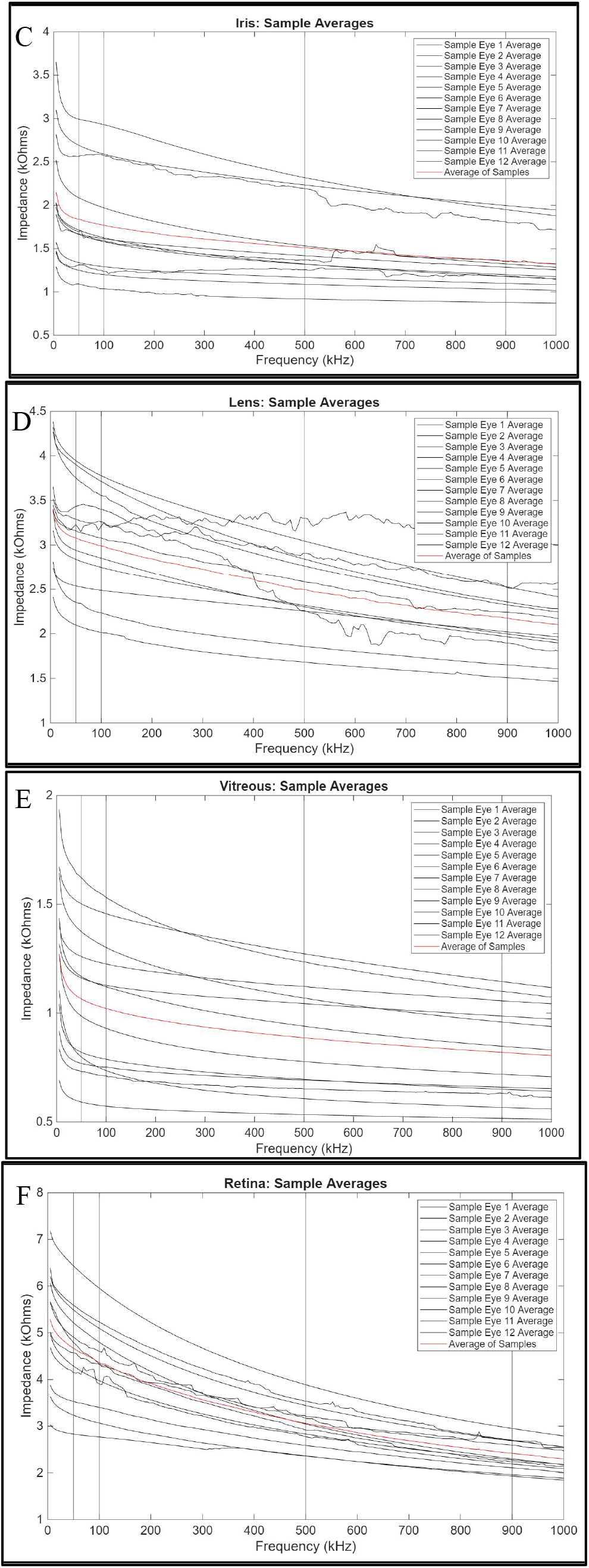
Average impedance (kΩ) of ocular tissues across a full frequency sweep (5 kHz–1 MHz): A) sclera, B) epithelial cornea, C) iris, D) lens, E) vitreous, and F) retina. These curves illustrate each tissue’s characteristic frequency-dependent behavior, with sample points highlighted at 5, 50, 100, and 900 kHz—frequencies evaluated for discrete-frequency analysis in Figure 5. While Figure 5 presents tissue differentiation at selected frequencies, Figure 6 provides the underlying continuous impedance profiles used to construct the tissue-specific bioimpedance library. Endothelial measurements (C) were acquired under open-sky/manual positioning due to robotic workspace limitations and were not included as features in the discrete-frequency analysis. Artifacts observed above ∼900 kHz in some retinal, lens, and iris traces are consistent with probe–tissue interface variability (e.g., tissue motion or contact pressure changes) and highlight the sensitivity of impedance measurements to interface conditions under practical acquisition scenarios.

Impedance values extracted at 5, 50, 100, and 900 kHz are summarized in Table 4 and shown as reference tissue signatures in Figure 7. Across all tissues, impedance magnitude decreased with increasing frequency, with a plateau behavior observed at higher frequencies. While mean impedance values differed across tissues, partial distributional overlap was present for specific tissue pairs at certain frequencies. Notably, despite this overlap, separability analyses (Tables 2 and 3) demonstrate that many tissue pairs remain statistically and practically distinguishable, whereas others exhibit moderate discrimination depending on frequency. These results emphasize that impedance-based tissue differentiation is governed by both absolute signal differences and distributional overlap, reinforcing the need for frequency-specific interpretation. Table 4 provides descriptive summary statistics (mean ± SD, and sample sizes) for each tissue at 5, 50, 100, and 900 kHz and serves as the reference library for feature selection. Tissue-to-tissue separability was evaluated using multiplicity-corrected pairwise comparisons and ROC-based analyses rather than one-sample tests (Table 2).

**Table 4:** Descriptive impedance summary statistics (mean ± SD, kΩ) and sample sizes for ocular tissues measured at 5, 50, 100, and 900 kHz under simulated surgical conditions. Values are provided as reference for feature selection and subsequent separability analyses.

| Tissue | n | 5 kHz (kΩ) | 50 kHz (kΩ) | 100 kHz (kΩ) | 900 kHz (kΩ) |
| --- | --- | --- | --- | --- | --- |
| Cornea | 12 | 68.73 ± 0.91 | 4.56 ± 1.90 | 3.93 ± 1.44 | 2.841 ± 0.756 |
| Iris | 12 | 2.68 ± 0.85 | 1.817 ± 1.001 | 1.686 ± 0.774 | 1.434 ± 0.466 |
| Lens | 12 | 3.54 ± 0.84 | 2.51 ± 0.448 | 2.36 ± 0.41 | 1.985 ± 0.276 |
| Retina | 12 | 4.91 ± 1.79 | 2.98 ± 0.949 | 2.73 ± 0.845 | 2.166 ± 0.622 |
| Sclera | 12 | 13.35 ± 7.09 | 3.01 ± 0.849 | 2.74 ± 0.73 | 2.212 ± 0.484 |
| Vitreous | 12 | 1.41 ± 0.37 | 0.89 ± 0.26 | 0.87 ± 0.25 | 0.808 ± 0.218 |

**Figure 7:**
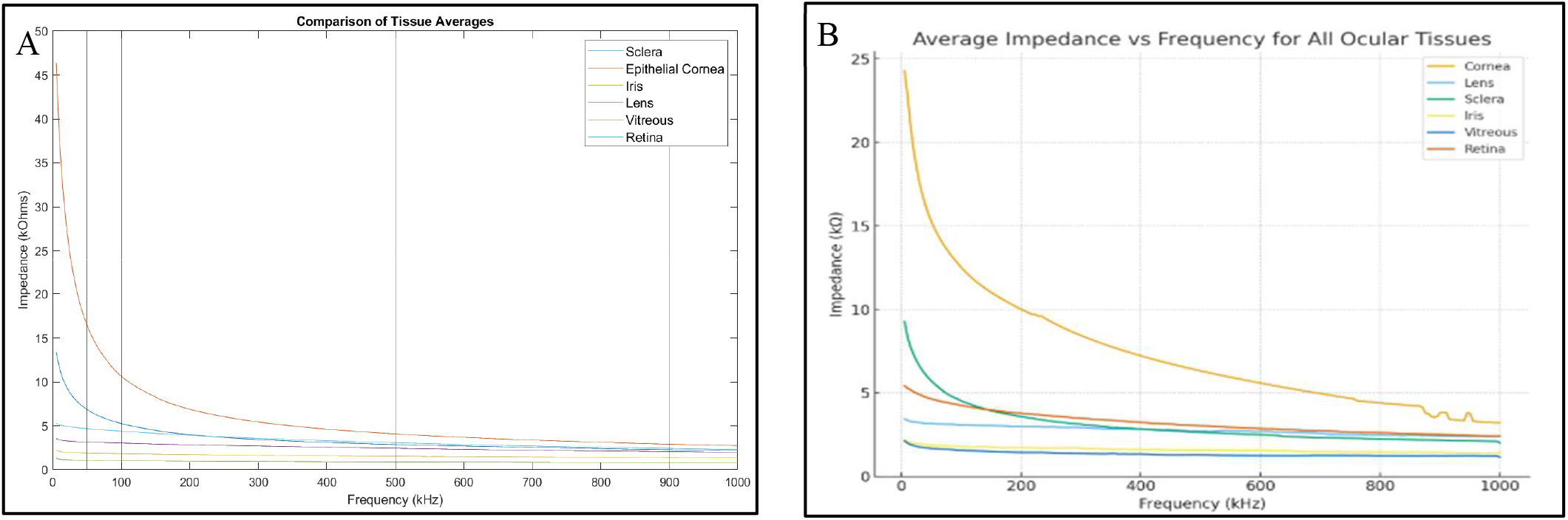
Tissue library constructed from impedance features sampled at 5, 50, 100, and 900 kHz. A) Reference tissue signatures derived from robotic-stabilized measurements. B) Handheld measurements acquired by an experienced ophthalmic surgeon. Agreement between robotic and handheld measurements is quantified using ICC and Bland–Altman analysis at the sampled frequencies (Table 5), demonstrating that handheld acquisition introduces increased variability relative to robotic stabilization while preserving the underlying tissue-specific impedance profiles.

Figure 7B shows impedance spectra acquired handheld by an experienced ophthalmic surgeon. Handheld acquisition introduces additional variability relative to robotic stabilization due to differences in probe positioning, contact force, and interface stability. To quantify this effect, robotic–handheld agreement was evaluated using intraclass correlation coefficients (ICC) and Bland–Altman analysis at the sampled discrete frequencies (Table 5). Agreement varied across tissues and frequencies, reflecting the sensitivity of impedance measurements to probe–tissue interface conditions under handheld acquisition. Negative or low ICC values observed in some tissue–frequency combinations reflect increased within-method variance associated with handheld probe positioning and interface variability rather than absence of tissue-specific impedance structure. This interpretation is supported by the preservation of tissue-specific impedance magnitude ranges and frequency-dependent trends observed in both robotic and handheld datasets (Table 4 and Figure 7).

**Table 5:** Agreement between robotic-stabilized and handheld impedance measurements at discrete frequencies. Intraclass correlation coefficients (ICC, two-way random effects, absolute agreement) and mean differences between robotic and handheld measurements are shown for each tissue at 5, 50, 100, and 900 kHz. ICC quantifies absolute agreement between acquisition modes, while mean differences quantify systematic bias. Agreement varied across tissues and frequencies, reflecting the sensitivity of impedance measurements to probe positioning, contact force, and probe–tissue interface stability during handheld acquisition. Full Bland–Altman limits of agreement are reported in Supplementary Table S3.

| Tissue | Frequency (kHz) | ICC (95% CI) | Mean difference (kΩ) |
| --- | --- | --- | --- |
| Cornea | 5 | -0.15 | -18.39 |
|  | 50 | 0.37 | 2.72 |
|  | 100 | 0.32 | 2.24 |
|  | 900 | -0.9 | 0.6 |
| Sclera | 5 | 0.23 | -3.51 |
|  | 50 | 0.61 | -0.35 |
|  | 100 | 0.6 | -0.29 |
|  | 900 | 0.59 | -0.16 |
| Iris | 5 | 0.1 | 2.32 |
|  | 50 | -0.29 | -0.49 |
|  | 100 | -0.34 | -0.33 |
|  | 900 | -0.4 | -0.08 |
| Lens | 5 | 0.66 | 0.24 |
|  | 50 | 0.46 | 0.15 |
|  | 100 | 0.47 | 0.24 |
|  | 900 | 0.43 | 0.34 |
| Vitreous | 5 | -0.13 | 0.94 |
|  | 50 | -0.23 | 0.52 |
|  | 100 | -0.23 | 0.51 |
|  | 900 | -0.22 | 0.47 |
| Retina | 5 | -0.13 | 0.86 |
|  | 50 | -0.54 | 0.18 |
|  | 100 | -0.66 | 0.21 |
|  | 900 | -0.78 | 0.27 |

Agreement between robotic-stabilized and handheld acquisition varied by tissue and frequency (Table 5). Bland–Altman analysis showed wider dispersion under handheld acquisition, consistent with greater variability in probe positioning and interface stability. Despite this increased variability, the overall impedance magnitude ranges and frequency-dependent trends observed in handheld measurements remained consistent with the robotic reference library (Table 4 and Figure 7). These findings support the use of robotic-stabilized measurements as the reference standard for library construction while defining the variability envelope associated with handheld acquisition.

Supplementary variability-controlled analyses demonstrated improved agreement between acquisition modes (Supplementary Table S3), indicating that probe–tissue interface stability is a major contributor to robotic–handheld dispersion. These findings indicate that impedance-based tissue differentiation in intact ocular environments is probabilistic rather than binary, with separability depending on frequency, tissue pair, and probe–tissue interface conditions.

## Discussion

This study establishes a physiologically relevant intact-eye ex vivo bioimpedance dataset for major ocular tissues measured under simulated surgical preparation. By preserving anatomical structure, tissue adjacency, and surrounding media, this work extends prior ocular bioimpedance studies performed on isolated tissues and captures boundary conditions that directly influence the electrical conduction pathway. The results demonstrate that ocular tissues exhibit reproducible, frequency-dependent impedance signatures that support the feasibility of impedance sensing as a complementary modality for intraoperative tissue identification.

Tissue differentiation was found to depend on both frequency and tissue pair. While global statistical differences were observed across tissues, practical separability varied across frequencies and between specific tissue contrasts. Mid-range frequencies provided the most consistent differentiation, whereas increased overlap was observed at lower and higher frequencies for certain tissue pairs. These findings indicate that no single frequency universally optimizes tissue discrimination and support the use of multi-frequency or frequency-tailored sensing strategies^14,15,17^.

The intact-eye experimental configuration represents an important advancement over prior studies using isolated tissues. Preservation of ocular geometry and surrounding media enables measurement of impedance under conditions that more closely approximate surgical environments. The observed impedance ranges and trends are consistent with prior reports of tissue-dependent electrical behavior and proximity-based impedance variation (e.g., Pedram et al. ^10^; Schoevaerdts et al. ^9^), while extending these findings to intact ocular systems under clinically relevant boundary conditions.

The descriptive impedance library provides insight into the observed separability behavior. At lower frequencies, tissues exhibit a broader dynamic range of impedance values, whereas at higher frequencies, impedance values decrease and partially converge. This frequency-dependent convergence contributes to increased distributional overlap for certain tissue pairs and highlights the importance of frequency selection when designing impedance-based sensing systems.

A key limitation of impedance sensing is its sensitivity to probe–tissue interface conditions, including contact pressure, micro-motion, tissue deformation, and hydration state ^18,20^. Variability observed in the frequency sweeps, particularly at higher frequencies, is consistent with interface-dependent effects and underscores a central challenge for translational implementation. Although probe–tissue contact was verified using OCT, contact force was not quantitatively controlled. As a result, the reported impedance signatures reflect controlled but not fully standardized interface conditions. These findings emphasize that interface stability is a dominant contributor to measurement variability and should be explicitly addressed in future system design ^18,20^.

The IRISS platform served as a stabilization mechanism to reduce operator-dependent variability and enable consistent probe placement during reference library construction ^11,12^. To evaluate translational applicability beyond robotic stabilization, handheld measurements acquired by an experienced ophthalmic surgeon were included as a variability stress-test. Agreement analysis using intraclass correlation coefficients and Bland–Altman metrics (Table 5) demonstrated tissue- and frequency-dependent agreement between robotic and handheld acquisition modes. Despite increased variability under handheld conditions, the preservation of tissue-specific impedance ranges and frequency-dependent trends indicates that underlying tissue signatures remain detectable under less controlled conditions.

The selected discrete frequencies span regions of the impedance spectrum where both contrast and stability are observed. In this dataset, mid-range frequencies provided improved overall separability across multiple tissue contrasts and therefore represent promising candidates for feature selection in future real-time implementations. However, frequency selection alone does not guarantee robust classification under variable contact conditions; therefore, future systems should incorporate decision thresholds and confidence measures validated against interface perturbations ^17,18^.

From a translational perspective, impedance sensing is best considered as a complementary intraoperative feedback modality rather than a standalone classifier. In scenarios where visual feedback is limited or depth perception is reduced, impedance measurements may provide real-time confirmation of tissue identity or proximity at the instrument tip. However, because distributional overlap exists for certain tissue pairs, particularly under variable interface conditions, practical deployment will likely require integration with additional sensing modalities or probabilistic decision frameworks to ensure reliability.

This study is limited by its ex vivo design, post-mortem sample variability, and lack of controlled intraocular pressure and physiologic perfusion. Post-mortem time effects within the sampled window are presented as exploratory and descriptive due to limited sample sizes per time bin; thus, we do not claim age-independence ^16^. Additionally, the current probe geometry targets major tissue classes encountered in cataract and vitreoretinal surgery and does not resolve microscale structures (e.g., trabecular meshwork and Schlemm’s canal). Translation toward in vivo validation will require expanded datasets with balanced sampling conditions, systematic characterization of probe–tissue interface mechanics, validation under physiologic intraocular conditions, and further probe miniaturization to improve spatial resolution.

Overall, this work provides an intact-eye ex vivo foundation for impedance-based tissue differentiation under surgically relevant conditions. By integrating statistical separability analysis with practical considerations of measurement variability, these results define both the capabilities and limitations of impedance sensing in ocular environments. With continued validation and development of interface-robust measurement strategies, bioimpedance sensing may contribute to future intraoperative feedback systems in robotic-assisted and handheld ophthalmic surgery ^9–12^.

## Conclusion

Intraocular microsurgery relies heavily on visual feedback, and the absence of direct tactile sensing—particularly in robotic-assisted workflows—limits the ability to detect unintended tissue contact at the instrument tip. This study establishes a physiologically relevant intact-eye ex vivo bioimpedance dataset for major ocular tissues measured under simulated surgical preparation, including BSS irrigation and OVD use, with probe–tissue contact verified using microscope visualization and OCT. Ocular tissues exhibited reproducible and frequency-dependent impedance differences, with separability varying by both tissue pair and sampling frequency. Mid-range frequencies provided the most consistent differentiation across tissues, whereas some tissue contrasts showed partial overlap depending on frequency and measurement conditions. Robotic stabilization reduced measurement variability and enabled construction of a reference impedance library, while handheld acquisition preserved tissue-specific trends within a broader variability envelope influenced by probe–tissue interface conditions.

Collectively, these findings provide a quantitative foundation for impedance-based ocular tissue differentiation and support the development of complementary intraoperative sensing strategies for ophthalmic microsurgery. Future work should focus on in vivo validation, improved control of probe–tissue interface mechanics, and integration with additional sensing modalities to support robust real-time deployment.

## Data repository

The datasets and classification code generated during this study are available from the corresponding author upon reasonable request.

## Acknowledgments

The authors acknowledge support from the UCLA Department of Mechanical and Aerospace Engineering, the NIH Vision T32 Training Fellowship (5T32EY007026), and NIH grant R01EY029689. The authors also used generative artificial intelligence (ChatGPT, OpenAI, GPT-5.3) in March 2026 for limited assistance with language editing and manuscript clarity. All scientific content and conclusions were developed and verified by the authors.

