## Supplementary Table S1 for "Frequency-Dependent Bioimpedance Signatures of Ocular Tissues in Intact Ex Vivo Eyes Under Simulated Surgical Conditions"

**Supplementary Table S1.** Pairwise statistical comparisons of impedance magnitude between ocular tissues at discrete frequencies (5, 50, 100, and 900 kHz).

| Frequency (KHz) | Tissue A | Tissue B | Mean Difference (kΩ) | Adjusted p-value | Hedges g | Cliff's δ |
|---|---|---|---|---|---|---|
| | Retina | Vitreous | 4.59 | 6.66E-14 | 3.83 | 1.00 |
| | Iris | Retina | -2.95 | 1.62E-04 | -2.26 | -0.82 |
| | Iris | Vitreous | 1.64 | 0.24 | 1.07 | 0.61 |
| | Cornea_epithelium | Iris | 3264.84 | 1.00 | 0.44 | 1.00 |
| | Cornea_epithelium | Lens | 3263.69 | 1.00 | 0.45 | 0.96 |
| | Cornea_epithelium | Retina | 3261.89 | 1.00 | 0.51 | 1.00 |
| | Cornea_epithelium | Sclera | 3143.72 | 1.00 | 0.38 | -0.05 |
| 5 | Cornea_epithelium | Vitreous | 3266.48 | 1.00 | 0.56 | 1.00 |
| | Iris | Lens | -1.14 | 1.00 | -0.34 | -0.21 |
| | Iris | Sclera | -121.12 | 1.00 | -0.70 | -0.99 |
| | Lens | Retina | -1.81 | 1.00 | -0.63 | -0.77 |
| | Lens | Sclera | -119.97 | 1.00 | -0.70 | -0.96 |
| | Lens | Vitreous | 2.78 | 1.00 | 0.99 | 0.70 |
| | Retina | Sclera | -118.17 | 1.00 | -0.79 | -0.99 |
| | Sclera | Vitreous | 122.76 | 1.00 | 0.89 | 1.00 |
| | Retina | Vitreous | 3.17 | 2.63E-13 | 3.11 | 0.97 |
| | Iris | Retina | -2.38 | 2.15E-07 | -2.59 | -0.90 |
| | Cornea_epithelium | Iris | 337.60 | 1.00 | 0.45 | 1.00 |
| | Cornea_epithelium | Lens | 335.75 | 1.00 | 0.46 | 0.90 |
| | Cornea_epithelium | Retina | 335.22 | 1.00 | 0.52 | 0.95 |
| | Cornea_epithelium | Sclera | 304.43 | 1.00 | 0.36 | 0.01 |
| | Cornea_epithelium | Vitreous | 338.39 | 1.00 | 0.57 | 1.00 |
| 50 | Iris | Lens | -1.85 | 1.00 | -0.59 | -0.62 |
| | Iris | Sclera | -33.17 | 1.00 | -1.23 | -0.98 |
| | Iris | Vitreous | 0.79 | 1.00 | 0.73 | 0.41 |
| | Lens | Retina | -0.53 | 1.00 | -0.19 | -0.69 |
| | Lens | Sclera | -31.32 | 1.00 | -1.18 | -0.92 |
| | Lens | Vitreous | 2.64 | 1.00 | 0.99 | 0.69 |
| | Retina | Sclera | -30.79 | 1.00 | -1.32 | -0.85 |
| | Sclera | Vitreous | 33.96 | 1.00 | 1.60 | 1.00 |
| | Retina | Vitreous | 2.78 | 1.13E-13 | 2.87 | 0.96 |
| | Iris | Retina | -2.14 | 1.77E-07 | -2.44 | -0.88 |
| | Cornea_epithelium | Iris | 171.05 | 1.00 | 0.46 | 0.98 |
| | Cornea_epithelium | Lens | 169.19 | 1.00 | 0.46 | 0.85 |
| | Cornea_epithelium | Retina | 168.91 | 1.00 | 0.52 | 0.75 |
| | Cornea_epithelium | Sclera | 148.43 | 1.00 | 0.35 | -0.05 |
| | Cornea_epithelium | Vitreous | 171.69 | 1.00 | 0.58 | 1.00 |
| 100 | Iris | Lens | -1.86 | 1.00 | -0.65 | -0.65 |
| | Iris | Sclera | -22.61 | 1.00 | -1.17 | -0.97 |
| | Iris | Vitreous | 0.64 | 1.00 | 0.65 | 0.40 |
| | Lens | Retina | -0.28 | 1.00 | -0.11 | -0.60 |
| | Lens | Sclera | -20.75 | 1.00 | -1.08 | -0.86 |
| | Lens | Vitreous | 2.51 | 1.00 | 1.02 | 0.70 |
| | Retina | Sclera | -20.48 | 1.00 | -1.22 | -0.72 |
| | Sclera | Vitreous | 23.26 | 1.00 | 1.52 | 1.00 |
| | Retina | Vitreous | 1.59 | 3.18E-13 | 2.77 | 0.93 |
| | Iris | Retina | -1.17 | 9.83E-07 | -2.27 | -0.85 |
| | Lens | Vitreous | 1.18 | 0.03 | 1.43 | 0.72 |
| | Sclera | Vitreous | 3.91 | 0.19 | 2.23 | 0.96 |
| | Iris | Sclera | -3.48 | 0.36 | -1.62 | -0.95 |
| | Iris | Lens | -0.75 | 1.00 | -0.86 | -0.45 |
| | Iris | Vitreous | 0.42 | 1.00 | 0.73 | 0.45 |
| 900 | Lens | Sclera | -2.73 | 1.00 | -1.21 | -0.71 |
| | Cornea_epithelium | Iris | 20.11 | 1.00 | 0.48 | 0.74 |
| | Cornea_epithelium | Lens | 19.36 | 1.00 | 0.47 | 0.59 |
| | Cornea_epithelium | Retina | 18.95 | 1.00 | 0.52 | 0.57 |
| | Cornea_epithelium | Sclera | 16.63 | 1.00 | 0.35 | 0.04 |
| | Cornea_epithelium | Vitreous | 20.54 | 1.00 | 0.62 | 0.88 |
| | Lens | Retina | -0.41 | 1.00 | -0.51 | -0.43 |
| | Retina | Sclera | -2.32 | 1.00 | -1.23 | -0.57 |

***Notes:*** *Adjusted p-values were computed using the Holm method following Games–Howell post-hoc comparisons. Hedges g is reported as a standardized effect size, and Cliff's δ provides a nonparametric measure of distributional separation. Positive values indicate higher impedance in Tissue A relative to Tissue B.*
