## Supplementary Table S2 for "Frequency-Dependent Bioimpedance Signatures of Ocular Tissues in Intact Ex Vivo Eyes Under Simulated Surgical Conditions"

**Supplementary Table S2.** Pairwise ROC–AUC, balanced accuracy, and optimal threshold-based classification between ocular tissues at discrete frequencies (5, 50, 100, and 900 kHz).

| Frequency (kHz) | Tissue A | Tissue B | n (Tissue A) | n (Tissue B) | ROC–AUC | Balanced Accuracy | Threshold (kΩ) | Decision Rule | Sensitivity | Specificity | AUC_sep |
|---|---|---|---|---|---|---|---|---|---|---|---|
| 5 | Cornea (epithelium) | Iris | 13 | 19 | 1.00 | 1.00 | 8.79 | predict Cornea (epithelium) if Z >= 8.793 kΩ | 1.00 | 1.00 | 1.00 |
| | Cornea (epithelium) | Vitreous | 13 | 40 | 1.00 | 1.00 | 7.31 | predict Cornea (epithelium) if Z >= 7.306 kΩ | 1.00 | 1.00 | 1.00 |
| | Sclera | Vitreous | 12 | 40 | 1.00 | 1.00 | 6.56 | predict Sclera if Z >= 6.565 kΩ | 1.00 | 1.00 | 1.00 |
| | Retina | Vitreous | 29 | 40 | 1.00 | 0.98 | 4.55 | predict Retina if Z >= 4.554 kΩ | 1.00 | 0.95 | 1.00 |
| | Iris | Sclera | 19 | 12 | 0.00 | 0.97 | 7.05 | predict Iris if Z <= 7.051 kΩ | 0.95 | 1.00 | 1.00 |
| | Retina | Sclera | 29 | 12 | 0.01 | 0.97 | 7.73 | predict Retina if Z <= 7.726 kΩ | 0.93 | 1.00 | 0.99 |
| | Cornea (epithelium) | Retina | 13 | 29 | 0.99 | 0.98 | 8.90 | predict Cornea (epithelium) if Z >= 8.903 kΩ | 1.00 | 0.97 | 0.99 |
| | Lens | Sclera | 19 | 12 | 0.02 | 0.97 | 7.85 | predict Lens if Z <= 7.853 kΩ | 0.95 | 1.00 | 0.98 |
| | Cornea (epithelium) | Lens | 13 | 19 | 0.98 | 0.97 | 8.59 | predict Cornea (epithelium) if Z >= 8.594 kΩ | 1.00 | 0.95 | 0.98 |
| | Iris | Retina | 19 | 29 | 0.08 | 0.95 | 4.55 | predict Iris if Z <= 4.554 kΩ | 0.89 | 1.00 | 0.92 |
| | Lens | Retina | 19 | 29 | 0.11 | 0.92 | 4.44 | predict Lens if Z <= 4.441 kΩ | 0.84 | 1.00 | 0.89 |
| | Lens | Vitreous | 19 | 40 | 0.84 | 0.85 | 2.08 | predict Lens if Z >= 2.075 kΩ | 1.00 | 0.70 | 0.84 |
| | Iris | Vitreous | 19 | 40 | 0.76 | 0.75 | 1.19 | predict Iris if Z >= 1.189 kΩ | 1.00 | 0.50 | 0.76 |
| | Iris | Lens | 19 | 19 | 0.37 | 0.71 | 2.05 | predict Iris if Z <= 2.049 kΩ | 0.42 | 1.00 | 0.63 |
| | Cornea (epithelium) | Sclera | 13 | 12 | 0.47 | 0.63 | 38.09 | predict Cornea (epithelium) if Z <= 38.09 kΩ | 0.77 | 0.50 | 0.53 |
| 50 | Cornea (epithelium) | Iris | 13 | 19 | 1.00 | 1.00 | 5.29 | predict Cornea (epithelium) if Z >= 5.291 kΩ | 1.00 | 1.00 | 1.00 |
| | Cornea (epithelium) | Vitreous | 13 | 40 | 1.00 | 1.00 | 4.71 | predict Cornea (epithelium) if Z >= 4.711 kΩ | 1.00 | 1.00 | 1.00 |
| | Sclera | Vitreous | 12 | 40 | 1.00 | 1.00 | 4.17 | predict Sclera if Z >= 4.167 kΩ | 1.00 | 1.00 | 1.00 |
| | Iris | Sclera | 19 | 12 | 0.01 | 0.97 | 3.92 | predict Iris if Z <= 3.923 kΩ | 0.95 | 1.00 | 0.99 |
| | Retina | Vitreous | 29 | 40 | 0.98 | 0.95 | 3.61 | predict Retina if Z >= 3.61 kΩ | 0.93 | 0.98 | 0.98 |
| | Cornea (epithelium) | Retina | 13 | 29 | 0.97 | 0.92 | 8.01 | predict Cornea (epithelium) if Z >= 8.01 kΩ | 0.85 | 1.00 | 0.97 |
| | Lens | Sclera | 19 | 12 | 0.04 | 0.95 | 4.12 | predict Lens if Z <= 4.123 kΩ | 0.89 | 1.00 | 0.96 |
| | Iris | Retina | 19 | 29 | 0.05 | 0.94 | 3.62 | predict Iris if Z <= 3.617 kΩ | 0.95 | 0.93 | 0.95 |
| | Cornea (epithelium) | Lens | 13 | 19 | 0.95 | 0.95 | 4.67 | predict Cornea (epithelium) if Z >= 4.666 kΩ | 1.00 | 0.89 | 0.95 |
| | Retina | Sclera | 29 | 12 | 0.07 | 0.92 | 6.66 | predict Retina if Z <= 6.663 kΩ | 1.00 | 0.83 | 0.93 |
| | Lens | Retina | 19 | 29 | 0.15 | 0.87 | 3.80 | predict Lens if Z <= 3.802 kΩ | 0.84 | 0.90 | 0.85 |
| | Lens | Vitreous | 19 | 40 | 0.83 | 0.80 | 1.76 | predict Lens if Z >= 1.761 kΩ | 1.00 | 0.60 | 0.83 |
| | Iris | Lens | 19 | 19 | 0.18 | 0.79 | 2.47 | predict Iris if Z <= 2.469 kΩ | 0.89 | 0.68 | 0.82 |
| | Iris | Vitreous | 19 | 40 | 0.65 | 0.76 | 1.21 | predict Iris if Z >= 1.211 kΩ | 1.00 | 0.53 | 0.65 |
| | Cornea (epithelium) | Sclera | 13 | 12 | 0.51 | 0.64 | 14.80 | predict Cornea (epithelium) if Z <= 14.8 kΩ | 0.69 | 0.58 | 0.51 |
| 100 | Cornea (epithelium) | Vitreous | 13 | 40 | 1.00 | 0.99 | 3.34 | predict Cornea (epithelium) if Z >= 3.343 kΩ | 1.00 | 0.98 | 1.00 |
| | Sclera | Vitreous | 12 | 40 | 1.00 | 0.99 | 3.44 | predict Sclera if Z >= 3.444 kΩ | 1.00 | 0.98 | 1.00 |
| | Cornea (epithelium) | Iris | 13 | 19 | 0.99 | 0.97 | 3.25 | predict Cornea (epithelium) if Z >= 3.249 kΩ | 1.00 | 0.95 | 0.99 |
| | Iris | Sclera | 19 | 12 | 0.01 | 0.97 | 3.35 | predict Iris if Z <= 3.35 kΩ | 0.95 | 1.00 | 0.99 |
| | Retina | Vitreous | 29 | 40 | 0.97 | 0.93 | 3.06 | predict Retina if Z >= 3.055 kΩ | 0.93 | 0.93 | 0.97 |
| | Iris | Retina | 19 | 29 | 0.05 | 0.94 | 3.13 | predict Iris if Z <= 3.132 kΩ | 0.95 | 0.93 | 0.95 |
| | Lens | Sclera | 19 | 12 | 0.07 | 0.89 | 3.32 | predict Lens if Z <= 3.318 kΩ | 0.79 | 1.00 | 0.93 |
| | Cornea (epithelium) | Lens | 13 | 19 | 0.93 | 0.89 | 3.22 | predict Cornea (epithelium) if Z >= 3.217 kΩ | 1.00 | 0.79 | 0.93 |
| | Cornea (epithelium) | Retina | 13 | 29 | 0.87 | 0.92 | 6.39 | predict Cornea (epithelium) if Z >= 6.39 kΩ | 0.85 | 1.00 | 0.87 |
| | Retina | Sclera | 29 | 12 | 0.14 | 0.88 | 6.29 | predict Retina if Z <= 6.285 kΩ | 1.00 | 0.75 | 0.86 |
| | Lens | Vitreous | 19 | 40 | 0.84 | 0.80 | 1.66 | predict Lens if Z >= 1.656 kΩ | 1.00 | 0.60 | 0.84 |
| | Iris | Lens | 19 | 19 | 0.17 | 0.82 | 2.00 | predict Iris if Z <= 1.999 kΩ | 0.84 | 0.79 | 0.83 |
| | Lens | Retina | 19 | 29 | 0.20 | 0.86 | 3.10 | predict Lens if Z <= 3.1 kΩ | 0.79 | 0.93 | 0.80 |
| | Iris | Vitreous | 19 | 40 | 0.64 | 0.76 | 1.15 | predict Iris if Z >= 1.15 kΩ | 1.00 | 0.53 | 0.64 |
| | Cornea (epithelium) | Sclera | 13 | 12 | 0.47 | 0.64 | 11.77 | predict Cornea (epithelium) if Z <= 11.77 kΩ | 0.69 | 0.58 | 0.53 |
| 900 | Iris | Sclera | 19 | 12 | 0.02 | 0.95 | 1.57 | predict Iris if Z <= 1.567 kΩ | 0.89 | 1.00 | 0.98 |
| | Sclera | Vitreous | 12 | 40 | 0.98 | 0.94 | 1.60 | predict Sclera if Z >= 1.6 kΩ | 1.00 | 0.88 | 0.98 |
| | Retina | Vitreous | 29 | 40 | 0.95 | 0.91 | 1.45 | predict Retina if Z >= 1.453 kΩ | 0.97 | 0.85 | 0.95 |
| | Cornea (epithelium) | Vitreous | 13 | 40 | 0.93 | 0.91 | 2.33 | predict Cornea (epithelium) if Z >= 2.329 kΩ | 0.85 | 0.98 | 0.93 |
| | Iris | Retina | 19 | 29 | 0.07 | 0.93 | 1.49 | predict Iris if Z <= 1.491 kΩ | 0.89 | 0.97 | 0.93 |
| | Cornea (epithelium) | Iris | 13 | 19 | 0.88 | 0.90 | 2.18 | predict Cornea (epithelium) if Z >= 2.184 kΩ | 0.85 | 0.95 | 0.88 |
| | Lens | Sclera | 19 | 12 | 0.14 | 0.82 | 2.90 | predict Lens if Z <= 2.902 kΩ | 0.89 | 0.75 | 0.86 |
| | Lens | Vitreous | 19 | 40 | 0.85 | 0.81 | 0.88 | predict Lens if Z >= 0.8758 kΩ | 1.00 | 0.63 | 0.85 |
| | Cornea (epithelium) | Lens | 13 | 19 | 0.79 | 0.82 | 4.25 | predict Cornea (epithelium) if Z >= 4.253 kΩ | 0.69 | 0.95 | 0.79 |
| | Retina | Sclera | 29 | 12 | 0.21 | 0.86 | 3.00 | predict Retina if Z <= 3 kΩ | 0.97 | 0.75 | 0.79 |
| | Cornea (epithelium) | Retina | 13 | 29 | 0.79 | 0.85 | 3.69 | predict Cornea (epithelium) if Z >= 3.686 kΩ | 0.69 | 1.00 | 0.79 |
| | Iris | Lens | 19 | 19 | 0.26 | 0.79 | 1.52 | predict Iris if Z <= 1.522 kΩ | 0.89 | 0.68 | 0.74 |
| | Lens | Retina | 19 | 29 | 0.29 | 0.73 | 1.99 | predict Lens if Z <= 1.992 kΩ | 0.63 | 0.83 | 0.71 |
| | Iris | Vitreous | 19 | 40 | 0.70 | 0.74 | 0.66 | predict Iris if Z >= 0.6552 kΩ | 1.00 | 0.48 | 0.70 |
| | Cornea (epithelium) | Sclera | 13 | 12 | 0.52 | 0.60 | 4.15 | predict Cornea (epithelium) if Z >= 4.155 kΩ | 0.69 | 0.50 | 0.52 |

***Abbreviations:*** *AUC, area under the curve; AUC_sep, direction-invariant separability metric defined as max (AUC, 1–AUC).*

***Notes:*** *Thresholds were selected to maximize balanced accuracy. Sensitivity and specificity are reported with respect to Tissue A.*
