## Supplementary Table S3 for "Frequency-Dependent Bioimpedance Signatures of Ocular Tissues in Intact Ex Vivo Eyes Under Simulated Surgical Conditions"

**Supplementary Table S3.** Agreement between robotic-stabilized and handheld impedance measurements across ocular tissues and frequencies. Intraclass correlation coefficients (ICC), Pearson correlation, and Bland–Altman statistics are reported.

| Tissue | Freq (kHz) | n | Robot Mean (kOhm) | Robot SD | Hand Mean (kOhm) | Hand SD (kOhm) | ICC(2,1) | Pearson r | Mean Difference (kOhm) | LoA low (kOhm) | LoA High (kOhm) |
|---|---|---|---|---|---|---|---|---|---|---|---|
| Sclera | 5 | 5 | 12.77 | 9.29 | 9.26 | 3.41 | 0.23 | 0.34 | -3.51 | -20.62 | 13.60 |
| | 50 | 5 | 3.10 | 1.11 | 2.75 | 1.07 | 0.61 | 0.60 | -0.35 | -2.28 | 1.57 |
| | 100 | 5 | 2.82 | 0.95 | 2.53 | 0.91 | 0.60 | 0.58 | -0.29 | -1.98 | 1.39 |
| | 900 | 5 | 2.26 | 0.63 | 2.10 | 0.68 | 0.59 | 0.56 | -0.16 | -1.36 | 1.05 |
| Cornea | 5 | 5 | 42.66 | 65.84 | 24.28 | 8.45 | -0.15 | -0.52 | -18.39 | -156.79 | 120.01 |
| | 50 | 5 | 4.12 | 2.60 | 6.84 | 3.42 | 0.37 | 0.49 | 2.72 | -3.40 | 8.83 |
| | 100 | 5 | 3.56 | 1.95 | 5.80 | 2.64 | 0.32 | 0.45 | 2.24 | -2.61 | 7.09 |
| | 900 | 5 | 2.63 | 1.00 | 3.24 | 1.70 | -0.90 | -0.78 | 0.60 | -4.42 | 5.62 |
| Iris | 5 | 5 | 61.20 | 132.06 | 2.65 | 1.26 | -0.01 | -0.33 | -58.54 | -318.20 | 201.11 |
| | 50 | 5 | 2.05 | 1.43 | 1.57 | 0.40 | -0.29 | -0.49 | -0.49 | -3.74 | 2.77 |
| | 100 | 5 | 1.85 | 1.05 | 1.53 | 0.34 | -0.34 | -0.49 | -0.33 | -2.79 | 2.13 |
| | 900 | 5 | 1.48 | 0.54 | 1.40 | 0.25 | -0.40 | -0.40 | -0.08 | -1.41 | 1.26 |
| Lens | 5 | 5 | 4.00 | 0.94 | 4.24 | 1.81 | 0.66 | 0.75 | 0.24 | -2.25 | 2.72 |
| | 50 | 5 | 2.72 | 0.36 | 2.87 | 0.25 | 0.46 | 0.49 | 0.15 | -0.48 | 0.78 |
| | 100 | 5 | 2.56 | 0.31 | 2.81 | 0.33 | 0.47 | 0.57 | 0.24 | -0.34 | 0.83 |
| | 900 | 5 | 2.08 | 0.18 | 2.42 | 0.36 | 0.43 | 0.91 | 0.34 | -0.08 | 0.77 |
| Vitreous | 5 | 5 | 1.40 | 0.38 | 2.34 | 1.50 | -0.13 | -0.30 | 0.94 | -2.31 | 4.19 |
| | 50 | 5 | 0.89 | 0.27 | 1.42 | 0.80 | -0.23 | -0.43 | 0.52 | -1.34 | 2.39 |
| | 100 | 5 | 0.86 | 0.26 | 1.37 | 0.76 | -0.23 | -0.44 | 0.51 | -1.27 | 2.29 |
| | 900 | 5 | 0.79 | 0.22 | 1.26 | 0.66 | -0.22 | -0.45 | 0.47 | -1.05 | 2.00 |
| Retina | 5 | 5 | 4.60 | 2.60 | 5.46 | 0.35 | -0.13 | -0.44 | 0.86 | -4.57 | 6.28 |
| | 50 | 5 | 2.86 | 1.45 | 3.04 | 0.74 | -0.54 | -0.49 | 0.18 | -3.59 | 3.95 |
| | 100 | 5 | 2.61 | 1.30 | 2.81 | 0.65 | -0.66 | -0.60 | 0.21 | -3.25 | 3.66 |
| | 900 | 5 | 2.08 | 0.97 | 2.35 | 0.51 | -0.78 | -0.71 | 0.27 | -2.43 | 2.97 |

***Abbreviations:*** *ICC, intraclass correlation coefficient (two-way random effects, absolute agreement); LoA, limits of agreement.*

***Notes:*** *Mean difference and limits of agreement were calculated using Bland–Altman analysis (robotic minus handheld measurements). Positive values indicate higher impedance measured under robotic stabilization.*
